# Microenvironment-specific modulation of macrophage function and tumour progression by ADAMTS1 through Syndecan-4 shedding

**DOI:** 10.64898/2025.12.03.690963

**Authors:** Silvia Redondo-García, Rita Caracuel-Peramos, Francisco Javier Rodríguez-Baena, Salvador Muñoz-Mira, Ana García-Muñoz, Raúl López-Domínguez, María del Carmen Plaza-Calonge, Belén López-Millán, Pedro Carmona-Sáez, Juan Carlos Rodríguez-Manzaneque

**Author notes:** Current address: Antibody and Vaccine Group, Centre for Cancer Immunology, School of Cancer Sciences, Faculty of Medicine, University of Southampton, Southampton General Hospital, SO16 6YD, Southampton, United Kingdom (Silvia Redondo-García). Current address: Instituto de Neurociencias CSIC-UMH, San Juan de Alicante, 03550 Spain.

## Abstract

Recent studies have emphasized the role of ADAMTS proteases in inflammation and immunity, particularly in the context of tumour progression. Since inflammatory cells can either constrain or promote tumour growth, understanding how ADAMTS proteases influence immune cell behaviour is crucial. Using syngeneic tumour models-B16F1 melanoma and Lewis Lung Carcinoma (LLC)-in *Adamts1* knockout (Ats1-KO) mice, we observed model-specific outcomes underscoring the complexity of ADAMTS1 function in cancer. Transcriptomic and functional analyses revealed broad alterations in the matrisome and immune-related pathways across both tumour types. Strikingly, while tumour progression was impaired in B16F1-derived tumours, the LLC model-characterized by a stronger myeloid component-showed no dependency on *Adamts1*. To investigate this apparent resistance, we experimentally depleted macrophages and uncovered a profound functional defect in this population in Ats1-KO mice. *In vitro* assays confirmed reduced macrophage phagocytic activity. Mechanistically, we identified the transmembrane heparan sulfate proteoglycan syndecan-4 (SDC4), a known ADAMTS1 substrate, as a key mediator of this activity. Together, these findings reveal a previously unrecognized ADAMTS1-SDC4 axis that links extracellular matrix remodelling to macrophage phagocytosis, ultimately shaping tumour cell clearance and tumour progression.

## Introduction

Tumour studies analysing the extracellular matrix (ECM) keep providing relevant clues to better understand the biology of oncological malignancies, their prognosis and treatments responsiveness [1–7]. Beyond the diversity of ECM components in each tissue context, ECM complexity is further shaped by the activity of extracellular remodelling proteases [8–15].

The study of this multifaceted scenario has inspired large-scale initiatives such as the Matrisome [16] and the Degradome [17], aiming to systematically define ECM components and their functional interactions. Certainly, the impact of an altered proteolysis in the microenvironment keeps being updated by studies on genetically modified animal models and in particular from cancer-related reports [18]. However, there is still much work to fully decipher the complexity around ECM dynamics. The heterogeneity inherent to human tumours remarks the necessity of a deeper knowledge of the crosstalk among all elements within the tumour microenvironment [19,20].

To date, studies of the extracellular protease ADAMTS1 (*<u>a</u> <u>d</u>isintegrin and <u>m</u>etalloprotease with thrombo<u>s</u>pondin motif 1*) showed its angiostatic and tumour blocking properties [21] but also pro-metastatic and tumorigenic activities [22–25]. Research using *Adamts1* knockout (Ats1-KO) mice revealed phenotypic consequences such as reduced body weight, kidney malformation, and impaired female fertility [26,27]. Additional reports have suggested roles in myocardial morphogenesis [28], aortic aneurysms [29], and immune regulation [30], similar to other members of its family [31]. Finally, the catalytic activity of ADAMTS1 has been reported on various proteoglycans [32–34] and further extracellular substrates [35–37].

Our work aimed to identify the mechanism of action of this protease using complementary *in vivo* models. Importantly, these studies disclosed different outputs, confirming model- and microenvironment-specific requirements. Beyond our previous work showing that stromal *Adamts1* deficiency alters immune responses in healthy and B16F1 tumour-bearing mice [30], the use of the LLC model uncovered additional functions.

The inherent properties of LLC tumour growth in presence or absence of stromal *Adamts1* were deeply assessed here, including immunohistology, flow cytometry and transcriptomics, guiding to the in vivo application of macrophage-depletion strategies. Surprisingly, closer examination of macrophages, particularly their phagocytic properties, revealed a previously unreported role of ADAMTS1 in modulating the macrophage phagocytic capacity, suggestively mediated by the transmembrane proteoglycan syndecan-4 (SDC4). Given the cell-surface location of this proteoglycan, and its decoration with sulfated heparan sulfate (HS) glycosaminoglycan (GAG) chains, it is plausible that SDC4 contributes to the fine regulation of “eat-me”/”don’t-eat-me” signals, as reported for SDC1 [38] and more broadly for the glycocalyx as a regulator of immune engagement [39]. Collectively, our findings underscore the importance of dissecting the extracellular landscape governing tumour progression, particularly the glycocalyx and its influence on phagocytic signalling. The discovery of this unanticipated immunoregulatory mechanism suggests new avenues to explore how ADAMTS proteases shape the tumour glycocalyx and, consequently, immune cell function.

## Results

### The absence of stromal *Adamts1* differently affects tumour progression and immune microenvironment in a lung cancer syngeneic model

As previously reported, our results showed the blockade of melanoma B16F1 tumours in the absence of stromal *Adamts1* (Ats1-KO) [25], in agreement with other studies using a spontaneous breast cancer model [24]. To further investigate the role of ADAMTS1 protease in tumour progression, now we extended our research to the well-established Lewis Lung Carcinoma (LLC) syngeneic model. For this, 2.5×10^5^ LLC cells were subcutaneously injected in WT and Ats1-KO C57Bl/6 mice, and tumour progression was monitored for 29 days. Surprisingly, unlike in the B16F1 model [25,30], our evaluation showed that the absence of stromal *Adamts1* did not alter LLC tumour growth (Figure 1A), neither the final weight compared to WT littermates (Figure 1B).

**Figure 1.**
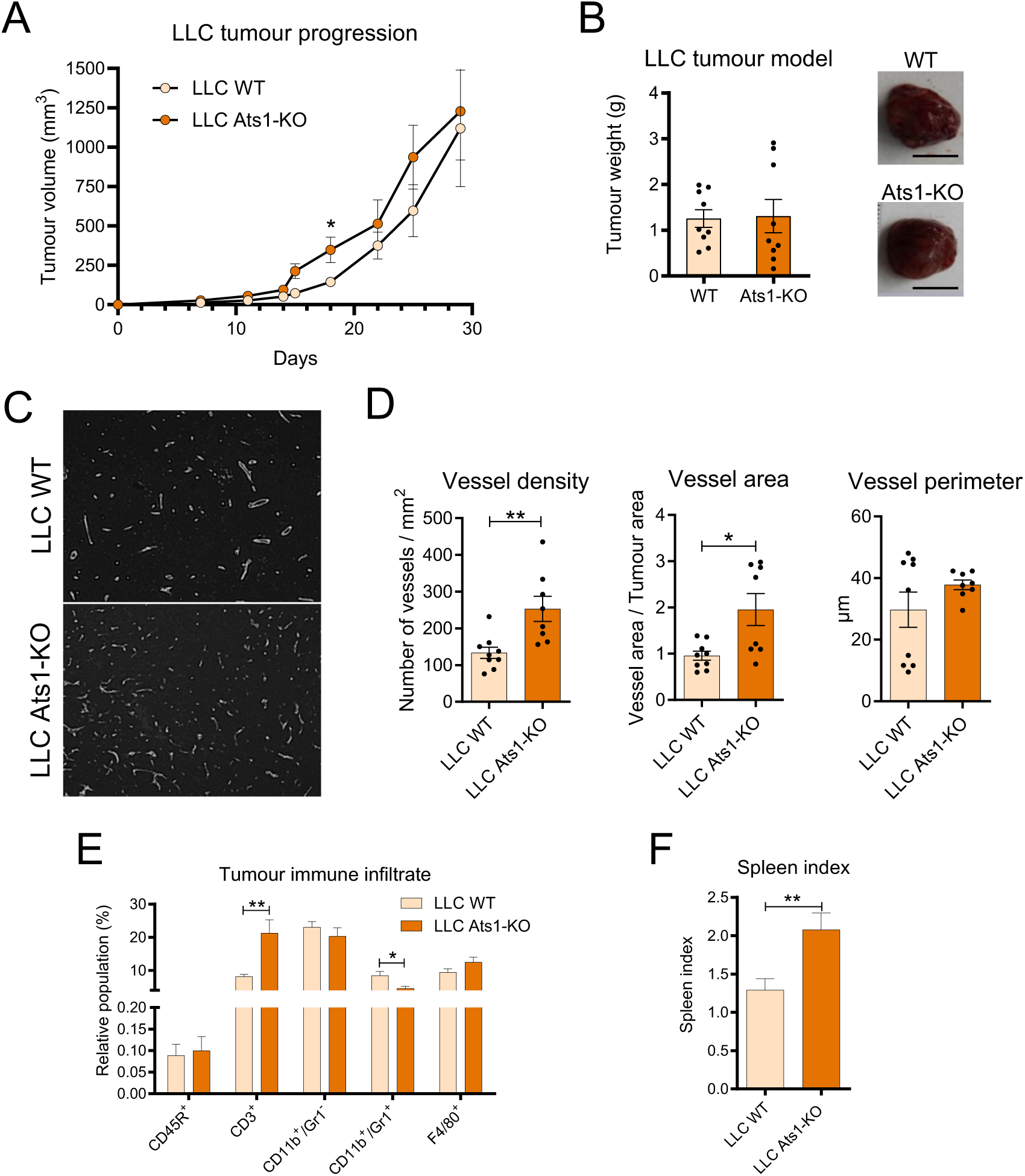
Effect of stromal ADAMTS1 on LLC tumour progression and immune landscape. (A) LLC tumour growth represented as tumour volume ± s.e.m. (B) Graphs representing mean LLC final tumour weight ± s.e.m, at day 29 in WT and Ats1-KO mice, and representative images of one tumour per group (black bar scale = 1 cm). (C) Fluorescence microscopy images of representative Endomucin-stained sections used for morphometric vasculature quantification in LLC WT and Ats1-KO tumours (white bar scale = 200 µm). (D) Graphs representing mean ± s.e.m of vessel density, vessel area and vessel perimeter from morphometric vasculature analysis of LLC WT and Ats1-KO tumours at endpoint. (E) FC data showing the percentage of infiltrated B cells, T cells, and myeloid cells (mean ± s.e.m of live cells) in LLC WT and LLC Ats1-KO tumours. (F) Graph showing spleen index ± s.e.m in LLC tumour-bearing WT and Ats1-KO mice. (*, *p*<0.05; **, *p*<0.01; two-tailed *t* Student; n=9 samples in LLC WT and LLC-Ats1-KO groups, respectively).

According to the recognized role of ADAMTS1 on angiogenesis, including our results in B16F1-derived tumours [25], we monitored and quantified the vasculature of LLC xenografts in WT and Ats1-KO mice using Endomucin staining (Figure 1C), as described [40]. Interestingly, vessel density and area were increased in Ats1-KO mice, although vessel perimeter remained unchanged (Figure 1D). However, further analysis of endothelial-related gene expression and immunofluorescence staining to assess vessel maturation revealed no significant differences (Supplementary Figure 1A-B). While the absence of stromal *Adamts1* led to some alterations in vasculature architecture, its lack of impact on tumour progression in the LLC model suggests the involvement of additional mechanisms.

Based on our previous findings, which demonstrated alterations in the bone marrow and spleen of healthy Ats1-KO mice, as well as an increase of CD3^+^ T cells within B16F1 tumours [30,41], we further analysed the immune infiltrate in LLC tumour-bearing mice (see Materials and Methods, and gating strategy in Supplementary Figure 2). Intriguingly, despite the absence of significant differences in tumour growth between groups (Figure 1A-B), we observed a similar increase in CD3^+^ T cells in Ats1-KO mice (Figure 1E), mirroring our observations in the B16F1 tumour model. In contrast, the effects on the myeloid compartment differed between the two models. In B16F1 tumours, various myeloid cell subpopulations were generally increased in Ats1-KO mice [30], whereas the LLC model only exhibited a decrease in CD11b^+^/Gr1^+^ cells in Ats1-KO tumours (Figure 1E). While these myeloid cells are typically associated with immunosuppressive functions, their reduction did not translate into detectable differences in tumour growth, suggesting a limited impact of stromal *Adamts1* deficiency in the LLC model compared to B16F1, as corroborated by gene expression analysis of immune-related genes (Supplementary Figure 3A-B). In addition, the overall population of F4/80^+^ macrophages remained unchanged in both tumour models, irrespective of *Adamts1* expression in the stroma (Figure 1E and [30]).

Given our previous findings on the immune-modulating effects of *Adamts1* deficiency in the spleen and bone marrow [30], we approached a more detailed analysis of these organs in LLC tumour-bearing mice. Despite the splenomegaly in Ats1-KO mice (Figure 1F), no significant differences were observed in any of the analysed immune cell populations in the spleen nor the bone marrow (Supplementary Figures 3C-D, respectively). These findings indicate that the loss of stromal *Adamts1* does not significantly alter immune cell proportions in distal organs in the LLC model, opposite to what was observed in both healthy and B16F1-bearing mice [30]. Moreover, this highlights the context-dependent effects of *Adamts1* on tumour progression, warranting further investigation into its underlying molecular interactions.

### RNAseq analysis reveals the impact of stromal *Adamts1* deficiency in tumour-specific matrisome and inflammatory signatures in LLC tumours

To better understand the tumour-specific roles of *Adamts1* and the distinct behaviour of both models in absence of this stromal protease, we first investigated the transcriptional landscape of LLC and B16F1 tumours by performing RNA sequencing in a WT background. The transcriptomic comparison revealed striking differences in their molecular profiles. Compared with LLC, B16F1 tumours displayed over 5000 upregulated genes, whereas the roughly 3000 genes downregulated relative to the LLC model included a prominent reduction of ECM-related transcripts such as *Col4a1, Lrp1, Col16a1*, *Col6a1, Ltbp1, Sema4c, Lama5, Lamb1, Col4a2, Mmp3, Sdc1,* and *Vcan* (Figure 2A), suggesting a more robust and transcriptionally active matrisome within the LLC tumour microenvironment.

**Figure 2.**
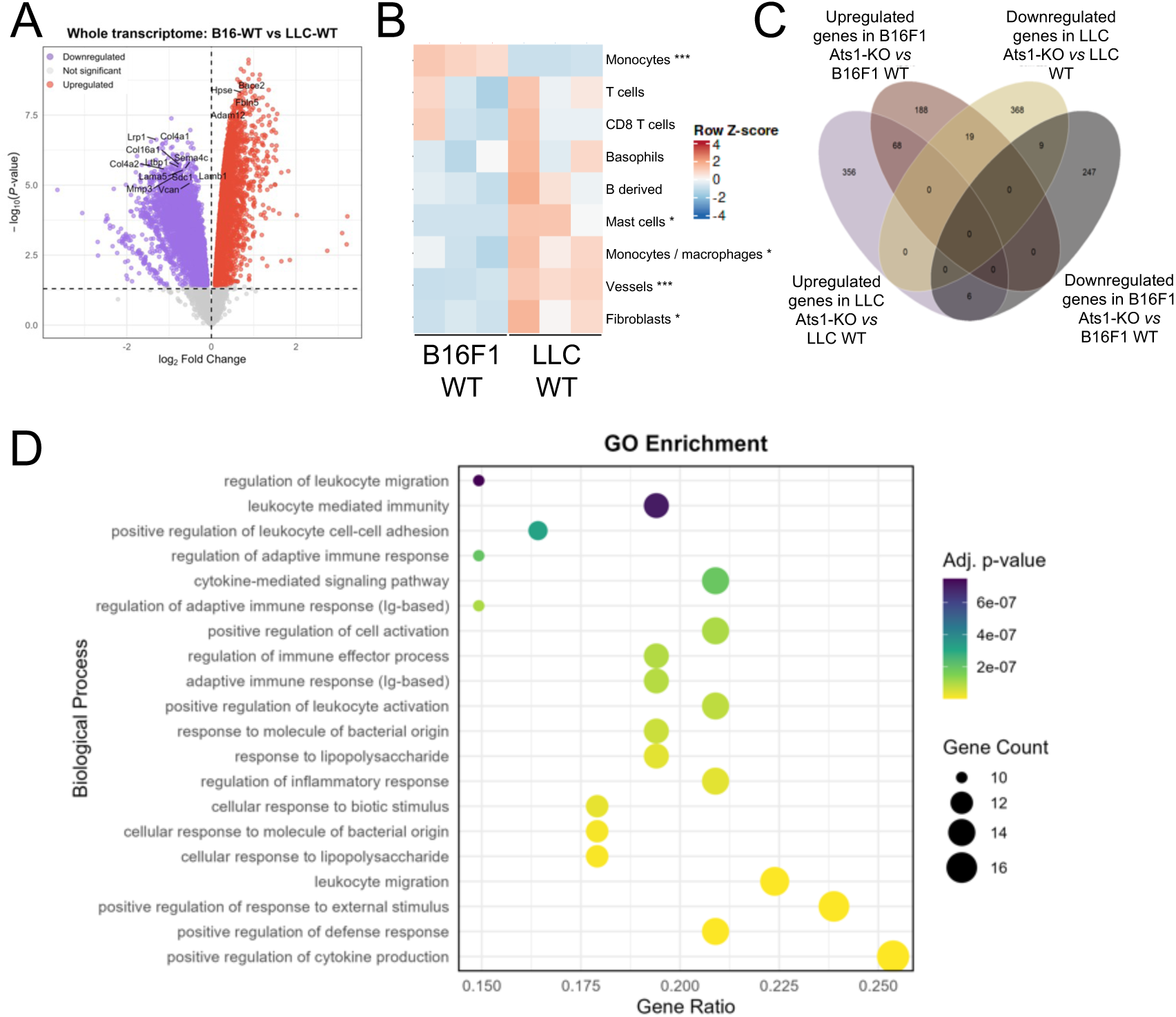
Whole transcriptome gene signature comparison between LLC and B16F1 tumours in WT and Ats1-KO conditions. (A) Volcano plot showing differential expression between B16-WT and LLC-WT tumours for the whole transcriptome. Each point represent a gene plotted as log_2_FC versus -log_10_ (adjusted *p* value). Genes significantly upregulated and downregulated genes in B16F1-WT relative to LLC-WT tumours are shown in red and purple, respectively; grey points denote non-significant genes. Highlighted genes correspond to ECM-related molecules. (B) Heatmap showing proportions of different cell types in LLC-WT and B16F1-WT tumours estimated with the mMCP-Counter software (*, *p< 0.05*; ***, *p<0.001*, t-test, n=3 per group). (C) Venn diagram depicting four sets of differentially expressed genes in LLC and B16F1 tumours in Ats1-KO versus WT mice. Overlapping areas indicate shared genes, while non-overlapping areas represent set-specific genes. (D) Gene Ontology (GO) enrichment analysis of the 68 genes significantly up-regulated in Ats1-KO tumours compared to WT counterparts for both LLC and B16F1 models. Gene ratio indicates the proportion of input genes associated with each biological process, while size and colour dot indicate the absolute number of genes (Count) and its significance, respectively. Biological processes are ranked by adjusted *p* value.

To further estimate the cellular composition underlying these transcriptomic differences, we applied mMCP-counter deconvolution analysis [42]. This analysis corroborated a more complex stromal compartment in LLC tumours, with higher inferred abundance of vasculature, fibroblasts and most immune populations.

Conversely, monocytes appeared less abundant in LLC, highlighting cell-type-specific differences in immunes landscapes between the two models (Figure 2B). To assess the specific impact of stromal *Adamts1* deficiency, we compared RNAseq profiles between WT and Ats1-KO tumours in both models. Across the whole transcriptome, 102 genes were differentially expressed in the absence of *Adamts1* in both models, which are mostly associated with Gene Ontology (GO) biological processes linked to inflammation and immune regulation. Interestingly, 68 genes were commonly upregulated in Ats1-KO tumours from both models (Figure 2C). These shared genes were enriched in GO biological processes related to inflammatory pathways, ranging from the strongest, which corresponded to leukocyte migration and immunity, to others related to cytokine production and signaling, immune effector processes, and adaptive immune responses (Figure 2D). These findings reinforce a direct association between *Adamts1* and inflammation, consistent with previous reports in the context of cachexia and in our earlier work [30,43], regardless of the tumour type. However, this shared inflammatory signature does not fully explain the different biological behaviour of LLC tumours compared with B16F1 in the absence of *Adamts1*, suggesting that additional, tumour-specific mechanisms must underlie the divergent phenotypes observed.

To understand these differences, we next focused on model-specific changes by examining the RNAseq profiles of LLC WT and LLC Ats1-KO tumours in greater depth. Whole-transcriptome analysis revealed significant alterations across multiple pathways, with a predominant enrichment in processes related to inflammation and immune response, including cytokine production, leukocyte modulation, and T cell activation (Figure 3A-B). These findings further underscore the strong impact of stromal *Adamts1* on immune-related pathways. A more focused examination of matrisome- and inflammation-associated signatures uncovered additional mechanistic insights. Analysis of the matrisome signature showed that loss of *Adamts1* (LLC Ats1-KO tumours) triggered a marked ECM remodelling response, affecting collagen organisation and broader ECM structural regulation (Figure 3C), which has been previously associated with pro-tumorigenic environments [44,45]. Remarkably, matrisome-associated differentially expressed genes also showed strong enrichment for pathways linked to cytokine production and signalling, leukocyte activation and migration, T cell differentiation, and phagocytosis, reinforcing the tight functional crosstalk between the extracellular compartment and the immune system. Within this context, matrisome-related genes upregulated in LLC WT tumours were predominantly associated with the positive regulation of defence responses, cytokine production, leukocyte activation, and myeloid cell migration (Figure 3D). Conversely, downregulated genes were linked to epithelial cell proliferation and ossification (Figure 3E). These results collectively highlight the crosstalk between the ECM and the immune response, and emphasize how the absence of *Adamts1* perturbs both compartments.

**Figure 3.**
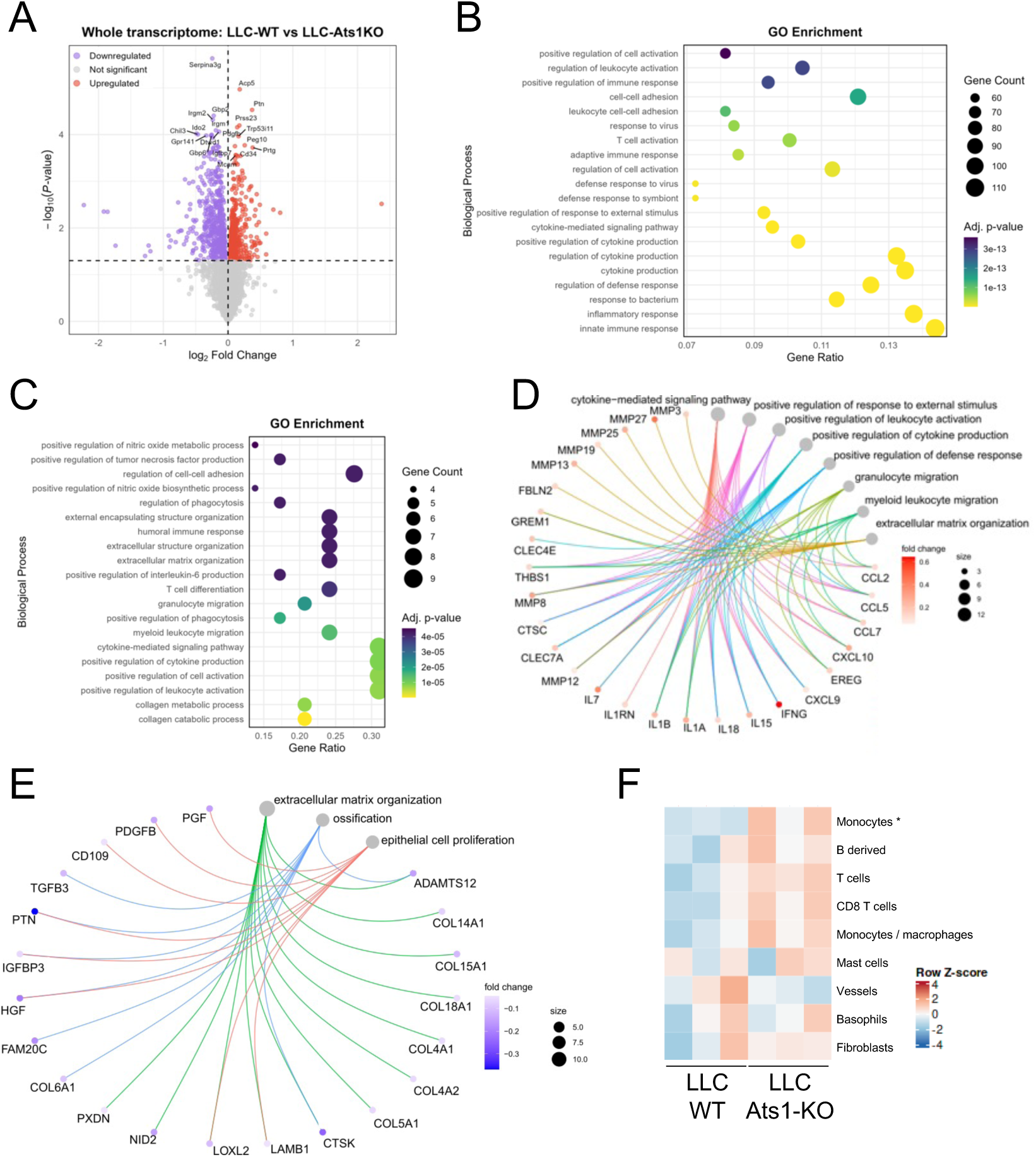
Impact of stromal *Adamts1* deficiency on matrisome- and inflammation-associated genes in LLC tumours. (A) Volcano plot showing the differential expression between LLC-WT and Ats1-KO tumours for the whole transcriptome. Each point represents a gene plotted as log_2_FC *versus* -log_10_ (adjusted *p* value). Genes significantly upregulated or downregulated in WT compared to Ats1-KO tumours are shown in red and purple, respectively; grey points denote non-significant genes. (B-C) GO enrichment of biological processes associated with differently expressed genes between LLC-WT and Ats1-KO tumours for the whole transcriptome (B) and for matrisome-related genes (C). The gene ratio represents the proportion of input genes associated with each biological process, while size and colour dot indicate the absolute number of genes (Count) and its significance, respectively. Biological processes are ranked by adjusted *p* value. (D-E) Network representation of genes involved in the most enriched GO biological processes (those with the highest significance and ≥ 8 genes involved) extracted from upregulated (D, red), and downregulated (E, purple) matrisome genes in the LLC WT versus LLC Ats1-KO comparison. More intense colours indicate higher fold change value. (F) Heatmap showing the proportions of different cell types in LLC WT and LLC Ats1-KO tumours estimated with the mMCP-Counter algorithm (*, *p< 0.05*, t-test, n=3 per group).

Finally, mMCP-counter deconvolution analysis revealed a higher inferred abundance of nearly all immune cell populations in LLC Ats1-KO tumours (Figure 3F), further demonstrating the broad impact of stromal *Adamts1* loss on the immune landscape.

Together, these findings indicate that *Adamts1* deficiency induces a coordinated inflammatory and ECM-remodelling response in LLC tumours. This response could explain why LLC tumour progression remains unaffected by the loss of *Adamts1*, in striking contrast to the dependency observed in the B16F1 model. Nonetheless, deeper analysis of specific immune cell subsets is necessary to determine whether *Adamts1* deficiency ultimately promotes a pro-tumorigenic immune niche in the LLC context.

### *In vivo* macrophage depletion reveals functional deficiencies of Ats1-KO macrophages

Macrophages play a critical role in tumour progression across various cancer types [46]. In particular, LLC tumours are characterized by a robust immune-myeloid compartment, which has been linked to therapy resistance [47]. Consistent with this, our cytometry analyses (Supplementary Figure 4) and mMCP-counter deconvolution (Figure 2B) revealed marked differences in myeloid infiltration between LLC and B16F1 tumours. Moreover, the matrisome-related transcriptomic alterations distinguishing LLC WT and LLC Ats1-KO tumours included pathways regulating myeloid migration and phagocytosis (Figure 3C), two core processes required for effective tumour clearance.

These observations, together with the limited impact of *Adamts1* deficiency on LLC tumour growth, prompted us to investigate the contribution of macrophages in vivo. We therefore depleted phagocytic myeloid cells using clodronate liposomes, a well-established method for macrophage ablation [48]. LLC cells were subcutaneously injected in WT and Ats1-KO mice, followed by treatment with control or clodronate liposomes (Figure 4A). Strikingly, tumour engraftment differed between genotypes. Whereas all Ats1-KO mice (5 out of 5) developed tumours following clodronate treatment, only 4 out of 6 WT mice exhibited tumour growth (Figure 4B). Tumour progression was also significantly delayed in clodronate-treated WT mice but remained unchanged in in Ats1-KO animals (Figure 4C). These findings indicate that the tumour-suppressive effect of clodronate liposomes is restricted to the WT scenario, suggesting impaired clodronate activity in Ats1-KO mice.

**Figure 4.**
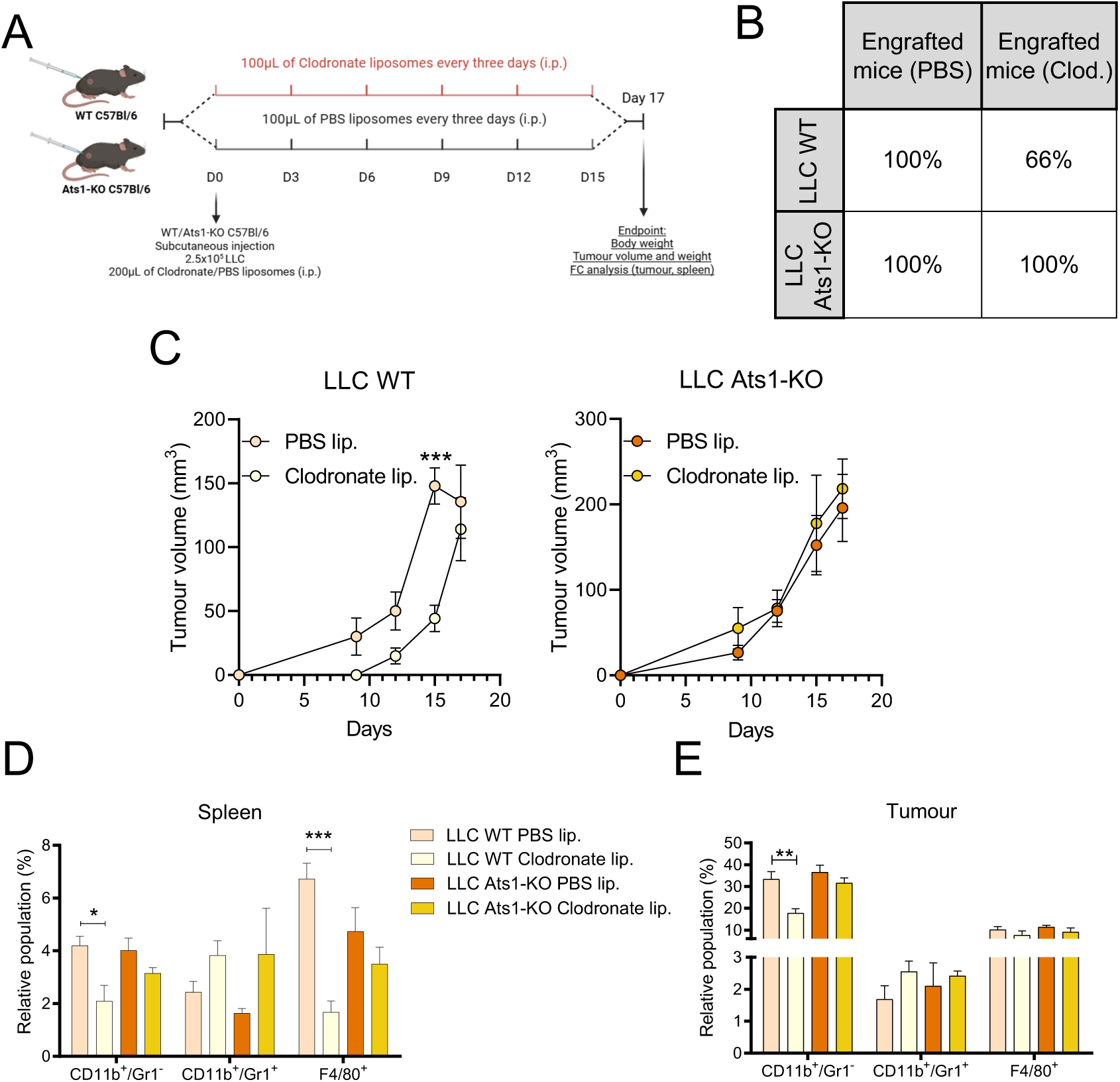
*In vivo* clodronate liposome treatment and analysis of myeloid depletion in LLC-bearing WT and Ats1-KO mice. (A) Experimental design for macrophage depletion in vivo using clodronate liposomes. (B) Table showing the percentage of tumour-engrafted mice in LLC WT and LLC Ats1-KO groups treated with PBS or clodronate liposomes. (C) Tumour growth curves (tumour volume ± s.e.m) for WT (left) and Ats1-KO (right) mice following PBS or clodronate liposome treatment. (D-E) FC analysis showing the percentage of myeloid populations (mean ± s.e.m) among live cells in spleens (D) and tumours (E) of WT and Ats1-KO mice treated with PBS or clodronate liposomes. (*, *p<0.05*; **, *p<0.01*; ***, *p<0.001*; two-tailed t Student, n=5, 4, 5 and 5 from left to right).

To assess the efficacy of myeloid depletion, we analysed spleen and tumour single-cell suspensions by FC. In WT mice, clodronate treatment led to a strong reduction of CD11b^+^/Gr1^-^ and F4/80^+^ macrophages (Figure 4D). In contrast, this effect was notably absent in Ats1-KO mice, indicating that myeloid depletion was markedly less effective in the *Adamts1*-deficient background. A similar trend was observed in tumours: CD11b^+^/Gr1^-^ cells were effectively reduced in WT but not Ats1-KO tumours following clodronate treatment, whereas F4/80^+^ macrophages showed minimal depletion in both genotypes, highlighting the heterogeneity of the tumour myeloid compartment (Figure 4E).

Because clodronate liposomes rely on phagocyte-mediated internalization to induce apoptosis [48], these results suggested that Ats1-KO macrophages may exhibit intrinsic functional impairments. To explore this, we generated bone marrow-derived macrophages (BMDMs) from WT and Ats1-KO mice and assessed their phenotypic and functional properties. While no major differences were observed in selection, morphology, adhesion or M1/M2 polarization (Supplementary Figure 5A-E), Ats1-KO macrophages displayed a mild, non-significant reduction in migratory capacity (Supplementary Figure 5F-G).

Together, these data indicate that *Adamts1* deficiency compromises the effectiveness of macrophage depletion, resulting in unchanged LLC tumour progression despite clodronate treatment. This impaired depletion suggests that Ats1-KO macrophages may harbour functional deficiencies, potentially affecting their behaviour within the tumour microenvironment. Whether these alterations contribute to a pro-tumorigenic immune landscape in the LLC model still requires further mechanistic investigation.

### In vitro assays confirm the deficient phagocytic activity of Ats1-KO macrophages

To determine whether *Adamts1* deficiency impairs macrophage phagocytic function, we performed a series of in vitro assays using BMDMs (Figure 5A and Supplementary Figure 6A). We first assessed tumour cell uptake by co-culturing BMDMs with CFSE-labelled LLC or B16F1 cells (Figure 5A, see Materials and Methods). Macrophages that phagocyted tumour cells were identified as CD11b^+^/CFSE^+^ cells by FC (Figure 5B). Across both tumour cell lines, the proportion of CD11b^+^/CFSE^+^ macrophages was consistently lower in Ats1-KO cultures compared with WT, demonstrating a clear reduction in tumour cell phagocytosis (Figure 5B-C). In parallel, in vitro exposure of BMDMs to clodronate liposomes showed a reduced apoptosis in Ats1-KO macrophages (Supplementary Figure 6B), consistent with the in vivo observation that clodronate-mediated depletion is inefficient in the *Adamts1*-deficient background and further supporting impaired phagocytic uptake.

**Figure 5.**
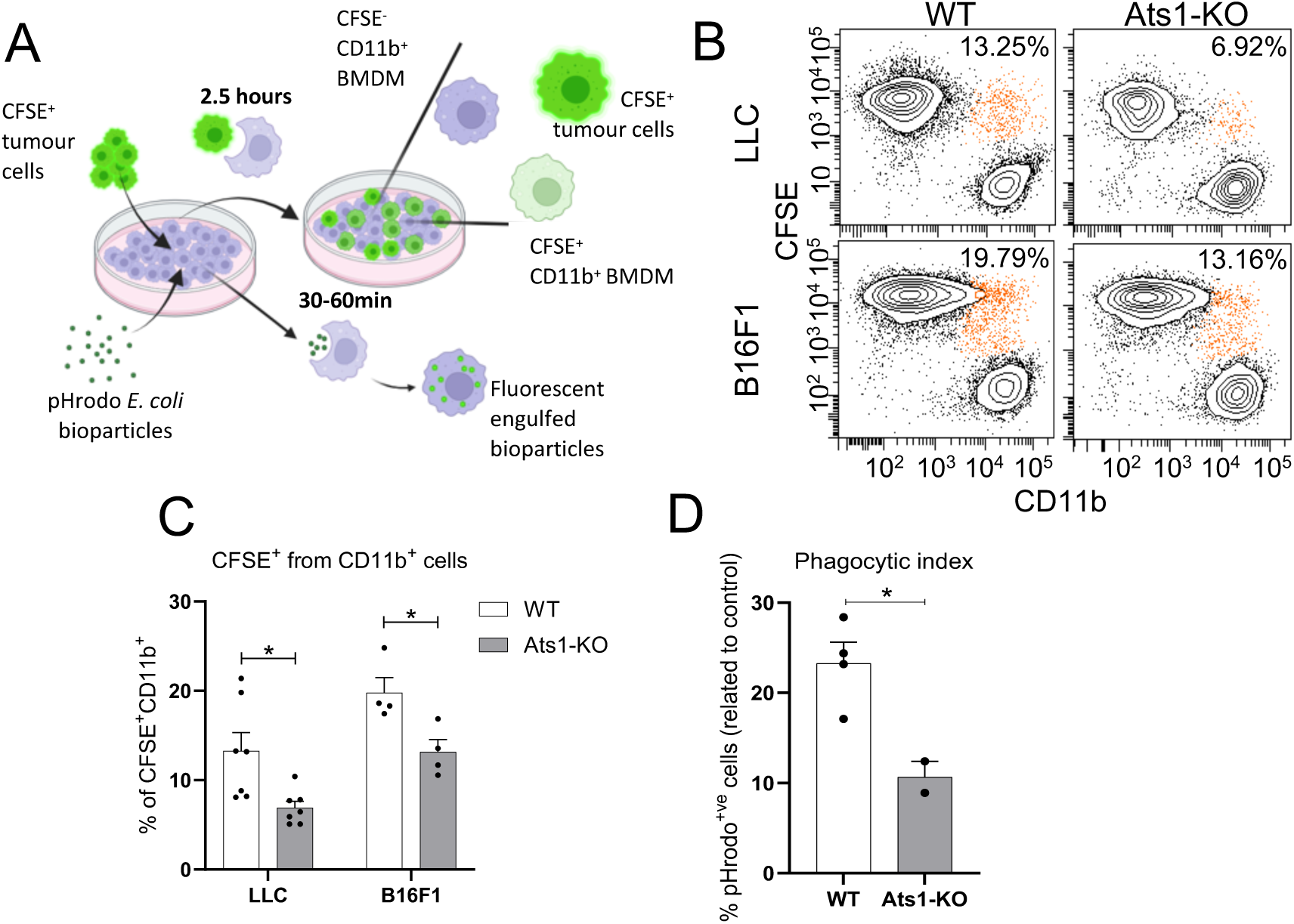
*In vitro* assessment of phagocytic activity in WT and Ats1-KO macrophages and the effect of metalloprotease inhibition. (A) Experimental design of phagocytosis assays based on co-culture of CFSE^+^-labelled tumour cells (top) and pHrodo^TM^-labelled *E.coli* bioparticles (bottom). (B) Representative FC plots showing the analysis of co-cultured tumour cells (CFSE^+^) and macrophages (CD11b^+^). (C) Quantification of CFSE^+^ cells within total CD11b^+^ macrophage populations (orange gate in panel B), showed as the mean percentage of live cells ± s.e.m (* *p<0.05;* two-tailed t Student, n=7 and 4 for LLC and B16F1 groups, respectively). (D) Phagocytosis assay using *E.coli* bioparticles in WT and Ats1-KO BMDMs, represented as the percentage of pHrodo positive cells (mean ± s.e.m) compared to the control (* *p<0.05;* two-tailed t Student, n=4, and 2 samples from WT and Ats1-KO groups, respectively).

To validate this defect using a different non-tumour-cell system, we performed phagocytosis assays with pHrodo^TM^-labelled, heat-inactivated *E.coli* bioparticles, which emit fluorescence upon engulfment (Figure 5A, see Materials and Methods). This assay also revealed a significant reduction in phagocytic activity in Ats1-KO BMDMs compared with WT controls (Figure 5D).

Collectively, these results demonstrate that Ats1-KO macrophages exhibit reduced phagocytic efficiency across multiple contexts, including tumour cell uptake, clodronate internalization, and bacterial particle engulfment. This deficiency unveils a previously unrecognized immunomodulatory function of ADAMTS1, likely mediated through its proteolytic processing of a cell-surface target. Given the pronounced myeloid infiltration in LLC tumours, impaired phagocytic activity in *Adamts1*-deficient macrophages may weaken anti-tumour immune responses in this model and help explain why LLC tumour progression remains largely unaffected by *Adamts1* loss.

### Shedding of SDC4, an ADAMTS1 substrate, as a mediator in macrophage phagocytic activity

Given the known proteolytic function of ADAMTS1, we next explored whether its enzymatic activity contributes to macrophage phagocytosis. WT BMDMs were treated with the broad-spectrum metalloprotease inhibitor BB94 and phagocytic activity was assessed using the pHrodo *E.coli* bioparticle assay. Although the reduction observed did not reach statistical significance, BB94 treatment reproduced the trend seen in Ats1-KO macrophages (Figure 6A), suggesting that metalloprotease activity contributes to optimal phagocytic function. Consistently, BB94 treatment also impaired *E. coli* uptake in RAW264.7 macrophage (Figure 6B).

**Figure 6.**
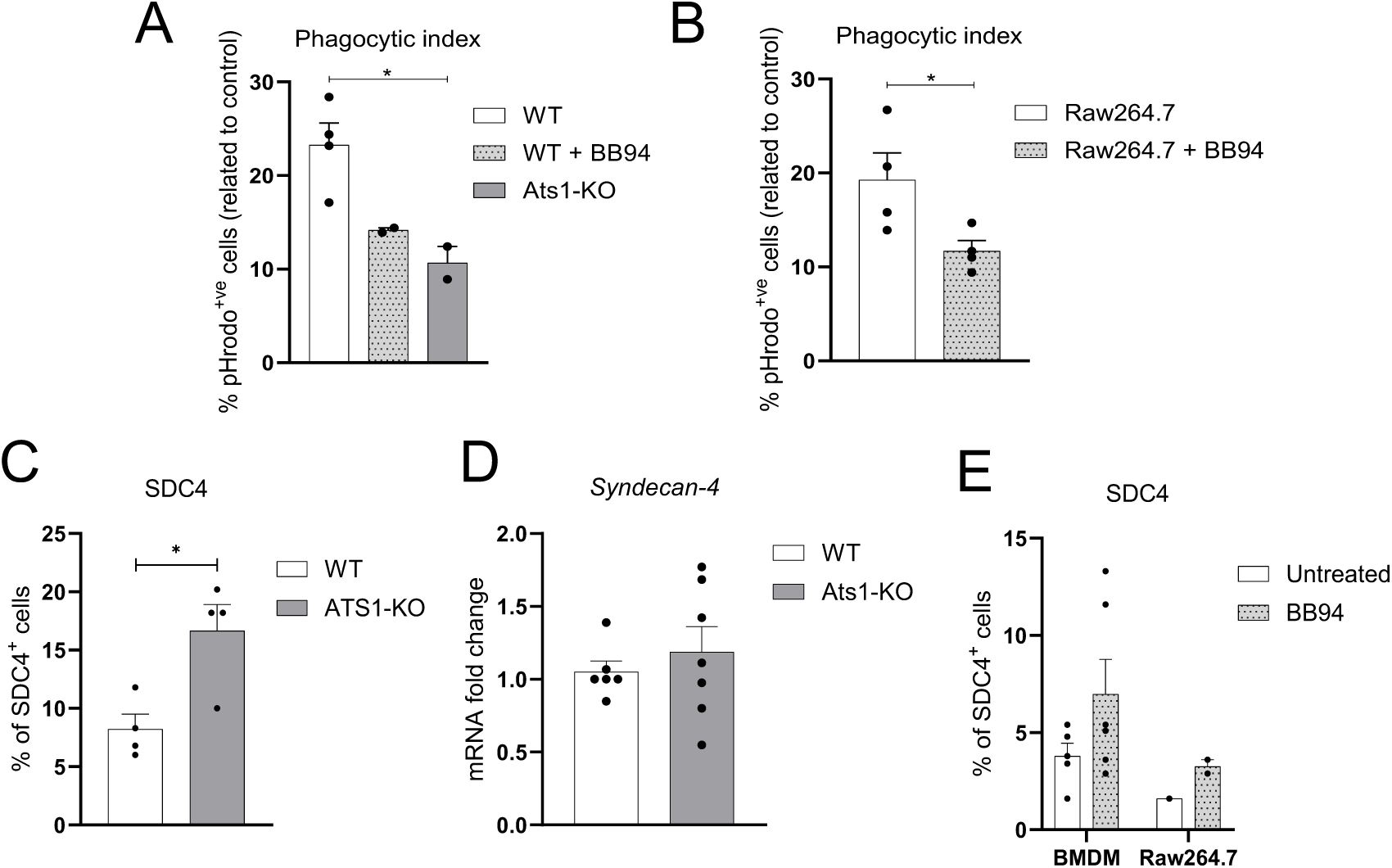
SDC4 and phagocytosis alterations in macrophages under protease inhibition or *Adamts1* deficiency. (A) Phagocytosis of pHrodo^TM^ *E,coli* bioparticles by WT BMDMs (with or without the metalloprotease inhibitor BB94), and Ats1-KO BMDM (mean ± s.e.m; * *p<0.05;* one-way ANOVA with multiple comparison; n=4, 2, and 2 from left to right, respectively). (B) Equivalent *E.coli* bioparticle uptake assay performed in RAW264.7 macrophages treated with or without BB94 (mean ± s.e.m; * *p<0.05;* two-tailed t Student; n=4 per group). (C) SDC4^+^ percentage from live cells and (D) *Sdc4* mRNA expression levels in WT and Ats1-KO BMDMs (mean ± s.e.m; * *p<0.05;* two-tailed t Student; n=4 per group). (E) Percentage of SDC4^+^ BMDMs and RAW264.7 cells treated or not with the metalloprotease inhibitor BB94 (mean ± s.e.m from live cells).

The macrophage glycocalyx, and particularly its heparan sulfate (HS)-rich components, has been implicated in modulating phagocytosis. Intact HS structures can sterically impede recognition of “eat-me” signals, thereby reducing target engulfment [38,49]. While HS-mediated regulation of inflammation is well-documented [50,51], including well-known HS-bearing transmembrane syndecans [38,52], still the functional consequences of their proteolytic shedding remain poorly understood.

Syndecan-4 (SDC4), a known substrate of ADAMTS1 [33], emerged as a strong candidate linking ADAMTS1 activity to the phagocytic dysfunction observed in *Adamts1*-deficient macrophages. ADAMTS1 cleaves the HS-modified ectodomain of SDC4, a process that could modulate cell-surface interactions critical for phagocytosis [38]. To test this, we quantified surface levels of the HS-bearing SDC4 ectodomain in WT and Ats1-KO BMDMs. Ats1-KO macrophages exhibited markedly elevated levels of SDC4 at the cell surface (Figure 6C), despite unchanged SDC4 mRNA expression (Figure 6D). This accumulation coincided with their reduced phagocytic activity (Figure 5B-D), consistent with previous reports showing that HS chains can mask phagocytic cues and restrict engulfment [38].

To further examine whether impaired proteolytic shedding alters SDC4 turnover and macrophage phagocytosis, we treated WT BMDMs and RAW264.7 cells with BB94. Inhibition of metalloprotease activity increased SDC4 surface levels (Figure 6E), correlating with a reduced phagocytosis (Figure 6A-B) and recapitulating the phenotype of Ats1-KO macrophages. Interestingly, BB94 also downregulated *Sdc1* and *Sdc4* transcripts, whereas *Adamts1* expression remained unchanged, suggesting compensatory transcriptional feedback upon sustained protease inhibition. Nonetheless, our data indicate that ADAMTS1 acts as the predominant sheddase for SDC4 under basal conditions, and that its absence disrupts surface remodelling of the macrophage required for efficient phagocytic function.

Together, these findings support a model in which ADAMTS1-mediated shedding of SDC4 alleviates inhibitory HS barriers and facilitates efficient macrophage phagocytosis (Figure 7). Under normal conditions, ADAMTS1 cleaves the HS-rich SDC4 ectodomain, enabling functional macrophage-target interactions. In contrast, loss of ADAMTS1 or inhibition of its proteolytic activity leads to the accumulation of intact SDC4 at the cell surface, increasing HS-mediated repulsion and thereby reducing macrophage engagement with tumour cells or bacterial particles. These mechanisms provides a new functional link between ADAMTS1 activity, glycocalyx remodelling, and innate immune competence.

**Figure 7.**
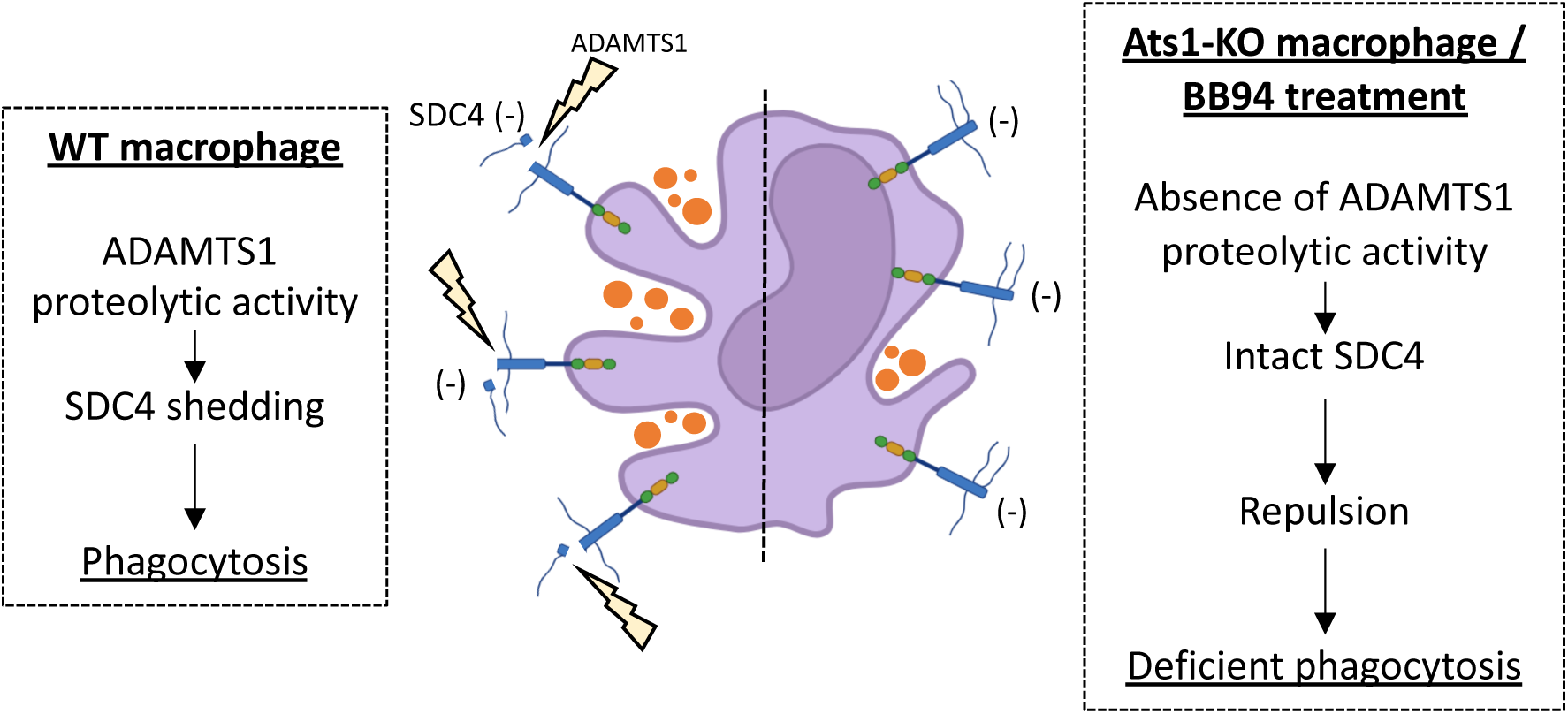
Schematic representation of SDC4 localization and its proteolytic shedding by ADAMTS1 on the surface of WT, Ats1-KO and BB94-treated macrophages, and its proposed mechanism of action during phagocytosis modulation.

## Discussion

Understanding the endogenous biological actions of extracellular proteases, such as ADAMTS1, has long been a major focus in the field, with particular emphasis in the identification of their substrates. Decades of research support the idea that physiological development depends on a finely tuned balance of extracellular molecules [53,54]; yet, despite this progress, our knowledge of this scenario remains incomplete.

Tumorigenesis, although often described as a chaotic process, still unfolds within the boundaries of conserved cellular and molecular mechanisms. Deepening our understanding of these mechanisms across distinct biological scenarios is therefore essential to fight pathologies and advance therapeutic strategies.

Reflecting the well-known heterogeneous nature of human cancer, our studies using different tumour models in WT and *Ats1*-KO mice revealed model-specific outcomes that will be important for translational interpretation. In the LLC model, we detected vascular alterations consistent with the angiostatic functions reported for ADAMTS1, but these changes did not significantly affect tumour growth. This parallels the well-documented resistance of LLC tumours to antiangiogenic therapies [47], suggesting that, in this setting, vascular modulation alone is insufficient to constrain tumour progression. Additionally, we confirmed that ADAMTS1 influences tumour immune infiltration, consistent with emerging links between extracellular proteases and immune regulation [31,55–57].

Our transcriptomic analyses further reinforced this view, revealing coordinated alterations in matrisome and inflammation-related gene signatures in Ats1-KO tumours. These findings point to a direct association between *Adamts1* and inflammatory pathways, in line with original studies in cachexia models and further work [30,43]. The convergence of ECM and inflammatory signals is particularly relevant in tumour settings, where extracellular context and immune surveillance are tightly intertwined.

Most notably, our efforts to deplete macrophages during LLC tumour development uncovered an unsuspected scenario: a direct role for ADAMTS1 in modulating macrophage phagocytic capacity, mediated by proteolytic processing of the extracellular domain of SDC4. Although ADAMTS proteases are increasingly recognized as immunomodulators, a direct mechanistic link between ADAMTS1 activity and macrophage phagocytosis has not been described previously. Shedding of transmembrane proteoglycans is known to regulate ECM homeostasis and cell-matrix interactions [58], and syndecans in particular, contribute to diverse physiological and pathological processes including wound healing and vascular biology [52,59]. Here, we uncover a novel function for SDC4 in macrophage phagocytosis and demonstrate that its shedding, rather than changes in transcription, serves as a rapid and local regulatory mechanism.

Phagocytosis is a complex, evolutionarily conserved process essential for tissue remodelling, host defence, and innate immune homeostasis, involving coordinated intracellular events and surface-exposed receptors and glycocalyx components [39,49]. In tumour biology, impaired clearance of dying tumour cells has been associated with disease progression and poor prognosis [60]. Consequently, therapeutic strategies aimed at enhancing tumour cell recognition by macrophages, such as CD47 blockade, have gained substantial interest [61]. Our findings add a new layer to this framework. The ADAMTS1-dependent shedding of SDC4 resembles the effect of classical agonists such as phorbol esters or cholesterol-depleting agents [33], raising intriguing questions about how these stimuli influence phagocytic processes more broadly. In endothelial cells, ADAMTS1-mediated SDC4 cleavage has been shown to affect cytoskeletal organisation [33], yet in macrophages the consequence appears to be an altered engulfment capacity. The downstream cytoskeletal mechanisms underlying this effect remain to be elucidated. Although SDC4 surface expression in macrophages is relatively low, our results demonstrate that minimal levels still are functionally meaningful, producing significant shifts in phagocytic activity. Supporting this notion, SDC4 has been reported to undergo constitutive shedding in human primary monocyte-derived macrophages [62], with functional implications for CXCL12/SDF-1 signalling. Our findings therefore extend the importance of SDC4 shedding to tumour-associated macrophage function and identify ADAMTS1 as a key regulator of this process.

## Materials and Methods

### Cell culture

Murine RAW264.7, B16F1 melanoma and Lewis Lung carcinoma (LLC) cells were cultured in DMEM supplemented with 10 % of fetal bovine serum (FBS) and 1 % of penicillin/streptomycin under standard conditions (37 °C, 5 % CO_2_ and 95 % relative humidity).

### Mouse colony handling and development of syngeneic tumours

C57BL/6 wild type and Ats1-KO mice [63] were maintained and bred at the *Centro de Investigación Biomédica-University of Granada* animal facility following approved guidelines. Genotyping was approached as described [30]. For the development of syngeneic tumours 2.5 x 10^5^ LLC cells (in 100 μl of Phosphate-Buffered Saline (PBS)) were subcutaneously injected in the right flank of C57Bl/6 WT or Ats1-KO mice. Animal status and tumour dimensions were monitored every 3 days after cell injection, up to 29 days, or until the tumour reached 1 cm in length. Final tumour volume was calculated *ex vivo* according to the formula: tumour volume = (π x length x width x height)/6 [22]. To evaluate splenomegaly, spleen index was determined according to the formula: *Spleen index = square root of spleen weight (x100) divided by body weight* [64]. All animals were handled and sacrificed following proper ethical guidelines and according to internationally accepted guidelines, approved by UGR Ethical Committee and Junta de Andalucía. Tumours were dissected and processed for further analysis, as indicated in other sections.

### Macrophage depletion in LLC tumour-bearing mice

Depletion of macrophages was approached treating LLC tumour-bearing mice with clodronate liposomes [48] (scheme in Figure 4A). Following the subcutaneous injection of LLC cells in the right flank, 200 µL of clodronate liposomes was intraperitoneally injected. After that, 100 µL of liposomes were injected five times every three days to prevent macrophage repopulation [65]. In every injection, PBS liposomes were inoculated as control of the treatment. At day 17 from cell inoculation, mice were sacrificed and macrophage depletion was evaluated by flow cytometry (FC) of spleen and tumours.

### Generation of bone marrow-derived macrophages and characterization

Monocytes were obtained from tibia and femur of both legs from C57BL/6 WT and Ats1-KO mice. Bones were flushed out with PBS and cell suspension was filtered, centrifuged, resuspended in Red Blood Cell (RBC) Lysis Buffer (150 mM NH4Cl, 10 mM KHCO3, 0.1 mM Na2 EDTA, pH 7.2-7.4) and incubated for 4 min at room temperature to remove red blood cells. Bone marrow cells were cultured at a concentration of 4 x 10^5^ cells/mL in 100 mm non-treated plates with DMEM low glucose with 20 % of heat-inactivated FBS, 2 % of P/S and 20 ng/mL of murine Macrophage Colony Stimulating Growth Factor (mM-CSF) (Peprotech, USA). Medium was refreshed at day 4 adding 4 mL of media and at day 7, non-attached cells were discarded and BMDM were detached with 4 mM EDTA. BMDM enrichment and purity was confirmed according to CD11b positivity by FC.

BMDM adhesion was evaluated using E-plates in the xCELLigence system (ACEA Bioscience). 4 x 10^4^ WT and Ats1-KO macrophages were cultured on Matrigel-coated wells (300 µg/mL). Impedance measurements were taken every 15 minutes for 4 hours.

BMDM migration was evaluated on 24-well 6.5 mm Transwell® with 8.0 µm Pore Polycarbonate Membrane Inserts (Corning). 5 x 10^4^ WT and Ats1-KO BMDM were cultured on the upper side of the membrane of Transwells. M0 medium (DMEM low glucose, 10 % of heat-inactivated FBS, 1 % of P/S, 20 ng/mL of M-CSF) with or without FBS was used for positive and negative control of the migration, respectively. After 24 hours, inserts were stained with 0.2 % crystal violet in a 20 % methanol solution. Migrated macrophages were visualized by plating the inserts on µ-Slide 8 well chambered coverslips (Ibidi) and 4-6 pictures of each membrane were taken on a PALM Microbeam Laser Microdissection system (Zeiss). For each WT and Ats1-KO group, average of migrated cells was calculated, and relative migration was computed comparing positive control versus negative control.

For *in vitro* BMDM polarization, 3 x 10^5^ cells/well were seeded in 24-well tissue plates and cultured in presence of M0 medium, M1 medium (M0 media supplemented with 20 ng/mL IFNɣ (Peprotech) and 100 ng/ml LPS (Sigma) for classically-activated macrophages or M2 medium (M0 media supplemented with 20 ng/ml IL-4 (Peprotech) for alternatively-activated macrophages. After 24 h, BMDM were detached and analysed by FC and RNA expression.

### Flow Cytometry analyses

Depending on the type of sample, different procedures were performed: spleens were disrupted physically using a 5 ml syringe as a pestle on a 70 μm cell strainer, while B16F1 and LLC tumours required a mechanical disaggregation process with scissors. Then, samples were incubated with 0,5% collagenase (C2799, Sigma-Aldrich) in PBS for 1 h at 37 °C, shaking every 10 min. After incubation, collagenase activity was blocked by adding DMEM supplemented with 10 % serum, cell suspension was flushed through a 19.5 G needle, filtered through a 70 μm cell strainer and spun 5 min at 300 g. Bone marrow cells were isolated as describe above. After getting single-cells suspensions, all samples were resuspended in RBC lysis buffer for 4 min at room temperature. Finally, cells were centrifuged and pellet was resuspended in FACS buffer (PBS 1X, 1 % FCS and 2 mM EDTA) for cell counting and antibody incubation.

For flow cytometry analyses of BMDM, BM, spleen or tumour cell suspension, samples were incubated with FcR blocking solution (2.4G2 anti-mouse CD16/CD32, 553142, BD Bioscience) in a 1 % BSA with 1 % FCS and 7-Amino-Actinomycin D (7-AAD, 00-6993-50, eBioscience) viability solution for 5 min at 4 °C, to reduce background and identify dead cells. Then, conjugated-primary antibody solution was added and incubated for 30 min on ice. After incubation, cells were spun at 300g for 5 min and finally resuspended in FACS buffer and analysed using a FACSCanto II (BD Bioscience, USA). In the case of intracellular NOS2 staining, cell fixation and permeabilization were performed after surface antigens staining following manufacturer protocol (Fix&Perm® Cell fixation and permeabilisation kit, GAS-002, Nordic MUbio, Netherlands). Antibodies used for FC were: hamster monoclonal anti-mouse CD3e-PE (12-0031, Thermofisher), rat anti-mouse/human CD45R/B220-FITC (110452, Thermofisher), rat anti-mouse CD11b-APC (553312, BD Bioscience), rat anti-mouse Gr-1-FITC (RB6-8C5, Miltenyi Biotech), rat anti-mouse F4/80-PE (123110, Biolegend), rat anti-mouse CD206-PE (141706, BD Bioscience), human monoclonal anti-mouse SDC4-PE (130-117-542, Miltenyi Biotec), and rat anti-mouse NOS2-PE (61-5920-80, eBioscience). The process of gating and population selection for spleen, BM and tumours is detailed in Supplementary Figure 2, which was performed following two antibodies panels: panel 1 including CD3e, CD45R and CD11b antibodies; and panel 2 including CD11b, Gr1 and F4/80 antibodies.

### Flow cytometry-based phagocytosis assays

Tumour cell phagocytosis by macrophages was performed adapting [66] protocol. Briefly, LLC or B16F1 tumour cells were labelled with the fluorescent dye carboxyfluorescein diacetate succinimidyl ester (CFSE). CFSE-labelled cells were then added to WT or Ats1-KO BMDMs (3 x 10^5^ cells per well, seeded 24 h earlier), at a BMDM:tumour cell ratio of 1:2 of in M0 medium. After 2.5 h of incubation, cells were harvested, stained for CD11b, and analysed by FC. Phagocytosis was quantified as the percentage of CFSE^+^/CD11b^+^ macrophages within the total CD11b+ population, using the formula: (CFSE^+^/CD11b^+^ cells / total CD11b^+^ cells) x 100, as previously described [66].

Phagocytosis of bacteria-derived particles was assessed using pHrodo™ Green *E. coli* Bioparticles Conjugate following manufacturer instructions (Invitrogen, P35366). BMDMs or RAW264.7 macrophages were pre-incubated on ice for 1 h in serum-free media. pHrodo *E.coli* particles were then added at a ratio of 1.5 µg per 100,000 cells (Figure 5A). Following particle addition, BMDMs and RAW264.7 cells were incubated at 37 °C for 60 min or 30 min, respectively. Phagocytosis was halted by placing plates on ice, and cells were collected for FC analysis. The phagocytosis index was calculated as the percentage of pHrodo™-positive macrophages normalized to cytochalasin-treated controls. When indicated, the metalloprotease inhibitor BB94 was added 16 h prior to initiating the phagocytosis assay.

### Apoptosis assay

Clodronate liposomes are widely used for the depletion of macrophages *in vivo* since its internalization provokes macrophage dead [48]. Following the same mechanism, clodronate liposomes can be used *in vitro* to evaluate macrophage phagocytic activity by measuring their apoptosis. With this purpose, 3 x 10^5^ BMDM from WT and Ats1-KO mice were seeded in M0 medium. The next day, M0 medium was replaced by new one containing PBS (1:2000 dilution) or clodronate (1:20000 dilution) liposomes. 24 h after adding the liposomes, BMDM were collected and analysed by FC for Annexin V (apoptosis marker) and 7-amino-actinomycin D (7AAD) solution (death marker) following manufacturer protocol (PE Annexin V Apoptosis Detection Kit I, 559763, BD Bioscience). Data was normalized to PBS-liposomes treated BMDM.

### Immunofluorescence

For the morphometric analysis of vasculature (vessel density, area and perimeter), tumour sections were subjected to immunofluorescence staining with a monoclonal rat anti-mouse Endomucin antibody (SC-65495, SCBT). Images were captured with the AxioImager A1 microscope (Zeiss), and converted to binary for further analysis with Image J software as indicated [25,40]. Additional immunofluorescence determinations included the incubation with anti-smooth muscle actin antibody (C6198, Sigma-Aldrich). For BMDM staining and characterization of their morphology, BMDM polarization protocol was performed on glass coverslips coated with 5 µg/cm^2^ fibronectin. After 24 hours polarization, cells were fixed in 4 % paraformaldehyde, permeabilized with 0.1 % Triton X-100 and stained for α-tubulin (sc-5546, SCBT) and Phalloidin-TRITC (Sigma-Aldrich) following standard protocols. Finally, nuclei were stained with 0.5 μg/mL 4’, 6-diamidine-2-fenilindol (DAPI). Fluorescence confocal images were captured with a LSM 710 confocal microscope (Zeiss).

### RNA isolation and quantitative RT-PCR

Total RNA was extracted from organs, tissue and tumour biopsies using the NucleoSpin RNAII kit (Macherery-Nagel). cDNA was synthesized with iScript cDNA Synthesis Kit (BioRad). qPCR reaction was performed in a 7900HT and QuantStudio 6 Flex Real-Time PCR systems (Applied Biosystems) using Fast SYBR green master mix (Applied Biosystems). qPCR data show the 2^-ΔΔCt^ value (indicated as mRNA fold change in all the graphs), using the 18S gene as housekeeping endogenous control. Values show mean ± standard error of the mean (s.e.m.). Primers used for these assays are indicated in Supplementary Table 1.

### RNA sequencing and data processing

Total RNA from B16F1 and LLC tumours generated in WT and Ats1-KO mice was isolated as previously described and quantity and purity was measured using Agilent 2100 BioAnalyzer (Agilent). Starting with 500-2000 ng of total RNA (RIN values > 8.6). Libraries were prepared using the TrueSeq Stranded mRNA Library Prep Kit (Illumina) according to manufacturer’s protocol, capturing poly-adenylated RNA by transcription by oligo-dT primer. Next, RNA was fragmented and cDNA was synthesized, 3’ends were adenylated, adapters and barcodes were ligated and it was enriched by PCR. mRNA library quality was evaluated, quantified and sequenced on NextSeq500 or NextSeq2000 platforms (Illumina). After quality checking, the fastq files were aligned to the mouse genome (GRCm38) using STAR v2.7.0a [67]. Gene expression values were quantified using RSEM v1.2.31 software [68]. Raw counts were filtered and normalised and batch effect was corrected. Differential expressed genes (DEGs) were obtained using the *edgeR* v3.40.2 [69] and *limma* v3.54.2 [70] packages in R v4.2.3 (R Core Team, 2023). Differential expression was calculated using log_2_ fold change (log_2_FC), with positive values indicating up-regulation and negative values indicating down-regulation. DEGs were considered statistically significant when the false discovery rate (FDR) corrected p-value < 0.05 to correct for multiple comparisons Values of adjusted *p*-value were calculated using the Benjamini-Hochberg (HB) method [71]. Downstream analyses of DEGs were conducted in R environment. Differential expression of genes was visualized by plotting volcano plots in *ggplot2* v4.0.0 [72]. Venn diagrams were generated using the *ggvenn* v0.1.10 [73] package to visualized and identify overlapping sets of differently expressed genes across LLC and B16 tumour models. DEGs related to the matrisome were identified using MatrisomeAnalyzeR v1.0.1 [74]. For DEGs associated with the inflammatory response, the <u>hallmark</u> was downloaded from the Gene Set Enrichment Analysis software (GSEA).

Functional enrichment analyses were conducted for DEGs (adj.p-value<0.05) of indicated comparisons. Biological processes for Gene Ontology (GO) annotations were identified using *clusterProfiler* v4.6.2 [75,76]. Networks of genes involved in relevant functional terms were visualized using *enrichplot* v1.18.4 [77].

Tumour immune microenvironment composition was explored using the MCP-counter v1.2.0 [42], a transcriptomic marker-based method to characterize the immune infiltration by estimating the relative abundance of specific cell types. Results were visualized by a heatmap using the *pheatmap* v1.0.13 R package [78] and significant differences between groups were assessed by conducting a t-test for each cell type.

### Statistical Analysis

All statistical analyses (excluding RNAseq analyses described above) were performed using GraphPad Prism 9 (GraphPad software Inc.). For quantitative RT-PCR, flow cytometry datasets, and spleen index measurements, statistical significance was assessed using either unpaired two-tailed Student’s t-tests or one-way ANOVA, according to each experimental design. Error bars represent the standard error of the mean (s.e.m.). Prior to statistical testing, distributions were examined for outliers using Tukey’s method.

## Data availability

All data generated or analysed during this study are publicly available in this article, in Supplementary Information files, and in Gene Expression Omnibus (GEO).

## Acknowledgments

The authors would like to thank members of JCRM’s laboratory and GENYO’s support units for helping with technical assistance and further discussion. In addition to stated financial support, this study was benefited from the collaborative work with COST Actions (INNOGLY CA18103, Mye-InfoBank CA20117 and IMMUNO-model CA21135).

This work has been supported by grants from Ministerio de Ciencia e Innovación from Spain, co-financed by FEDER (PID2019-104416RB-I00 to JCRM), from Consejería de Salud y Familias Junta de Andalucía (PE-0225-2018), and from Consejería de Universidad, Investigación e Innovación Junta de Andalucía (PROYEXCEL_00877). SMM was funded by Primera Experiencia Program (SC/PEX/0086/2022), and AGM by Investigo Program (GR/INV/0013/2022).

## Author contributions

SRG, RCP, FJRM, and SMM performed experiments; MBLM contributed with the interpretation of FC data; AGM, RLD and PC contributed with bioinformatic studies; MCPC contributed with animal care, cell culture, and immunohistochemistry techniques; SRG, RCP, and JCRM conceived and designed the experiments; SRG and JCRM wrote the manuscript; all the authors read and contributed on the final edition of the manuscript; JCRM supervised the study.

