## Supplementary Information for "Microenvironment-specific modulation of macrophage function and tumour progression by ADAMTS1 through Syndecan-4 shedding"

**Silvia Redondo-García et al.**

**Supplementary Table 1.** Primers used for quantitative RT-PCR

| Gene | Specie | Forward | Reverse |
|---|---|---|---|
| ***Cdh5*** | Mouse | TTACTCAATCCACATACACATTTTCG | GCATGATGCTGTACTTGGTCATC |
| ***Cd31*** | Mouse | TCCAGGTGTGCGAAATGCT | TGGCAGCTGATGCCTATGG |
| ***Endoglin*** | Mouse | TCGATAGCAGCACTGGATGAC | AGCTTCTGGCAAGCACAAGAA |
| ***Cd3*** | Mouse | CAGTCAAGAGCTTCAGACAAG | GATGGCTGTACTGGTCATATTC |
| ***Cd4*** | Mouse | GAGTTCCCAGAAGAAGATCAC | AAGGCGAACCTCCTCTAA |
| ***Cd8*** | Mouse | CCATGAGGGACACGAATAATAA | GAGTTCACTTTCTGAAGGACTG |
| ***Cd11b*** | Mouse | GCAGCACTGAGATCCTGTTTA | CTCCACTTTGGTCTCTGTCTTAG |
| ***Cd163*** | Mouse | GAGGAAACTGTAAGTCGCTGAA | ACGGCACTCTTGGTTTGT |
| ***Cd206*** | Mouse | TATGGCAACAGACAAGAGAAG | GGAGTACATGGCTTCATATCC |
| ***FoxP3*** | Mouse | CAATAGTTCCTTCCCAGAGTTC | TCGGATAAGGGTGGCATAG |
| ***Nos2*** | Mouse | CTTGGTGAAAGTGGTGTTCTTTG | TCAGACTTCCCTGTCTCAGTAG |
| ***Sdc4*** | Mouse | ATGCTGGCGGCTCGGATGACT | GGGCTCAATCACTTCAGGGAAG |
| ***Tnfα*** | Mouse | GCCTCCCTCTCATCAGTTCTAT | CACTTGGTGGTGGTTTGCTACGA |
| ***18S*** | Mouse | CGGACAGGATTGACAGATTG | CAAATCGCTCCACCAACTAA |

**Supplementary Figure 1. Analysis of endothelial-related molecules in LLC tumours.** (A) Graph representing relative mRNA fold change expression ± s.e.m of vasculature-related genes (*Endoglin*, *Cdh5* and *Cd31*) in LLC tumours (n=9 per group). (B) Representative confocal microscopy images showing smooth muscle actin deposition (white) together with the endothelial marker Endomucin (red) in LLC tumours at endpoint (white bar scale = 50µm).

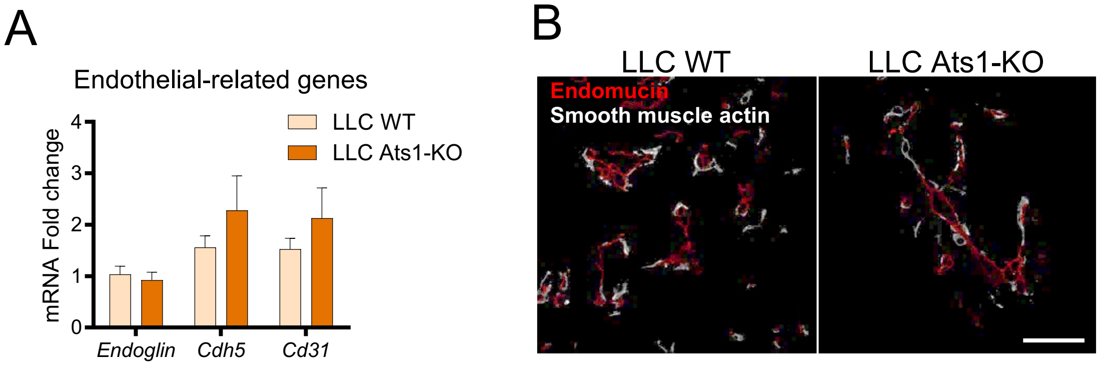

**Supplementary Figure 2. FC gating strategy and immune population identification.** Representative FC plots from spleen (A), bone marrow (B), and LLC tumours (C). Total cell populations were selected, and dead cells ($7AAD^{+}$) were excluded from subsequent analyses. Cells were stained using two antibody panels: Panel 1 (CD3-PE, CD45R-FITC, and CD11b-APC) and Panel 2 (CD11b-APC, Gr1-FITC, and F4/80-PE). $CD3^{+}$ T cells and $CD45R^{+}$ B cells were quantified from Panel 1 (top), while $CD11b^{+}/Gr1^{-}$ myeloid cells, $CD11b^{+}/Gr1^{+}$ myeloid-derived suppressor cells, and $F4/80^{+}$ macrophages were quantified from Panel 2 (bottom), all represented in black.

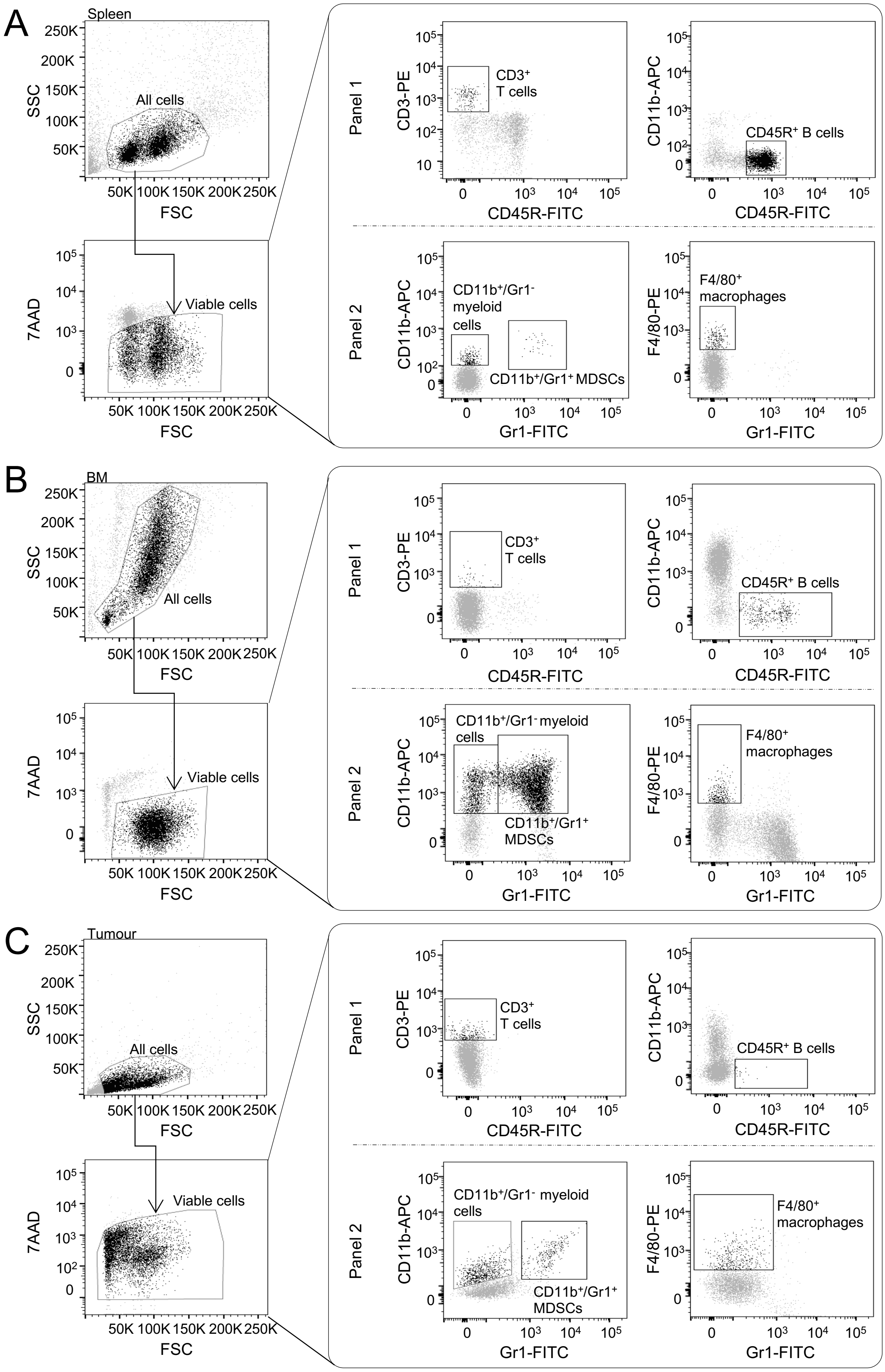

A
Spleen
All cells
Viable cells
B
BM
All cells
Viable cells
C
Tumour
All cells
Viable cells
SSC
FSC
7AAD
Panel 1
Panel 2
CD3-PE
CD45R-FITC
CD3+ T cells
CD11b-APC
CD45R+ B cells
Gr1-FITC
CD11b+/Gr1- myeloid cells
CD11b+/Gr1+ MDSCs
F4/80-PE
F4/80+ macrophages

**Supplementary Figure 3. Characterization of immune-related genes and populations in LLC and B16F1-tumour bearing WT and Ats1-KO mice.** (A, B) Graphs representing relative mRNA fold change expression ± s.e.m of genes related to T cells (*Cd3*, *Cd4*, *Foxp3* and *Cd8*, left) and macrophages (*Cd11b*, *Nos2*, *Tnfa*, *Cd163* and *Cd206*, right) of LLC (A) and B16F1 (B) tumours. (C, D) FC data showing the percentage of immune populations represented as mean ± s.e.m of live cells in WT and Ats1-KO spleens (C) and bone marrow (D) in WT and Ats1-KO LLC tumour-bearing mice (*, $p<0.05$; two-tailed t Student, n=9 and n=8 samples in LLC model and n=5 and n=6 samples in B16F1 model in WT and Ats1-KO groups, respectively).

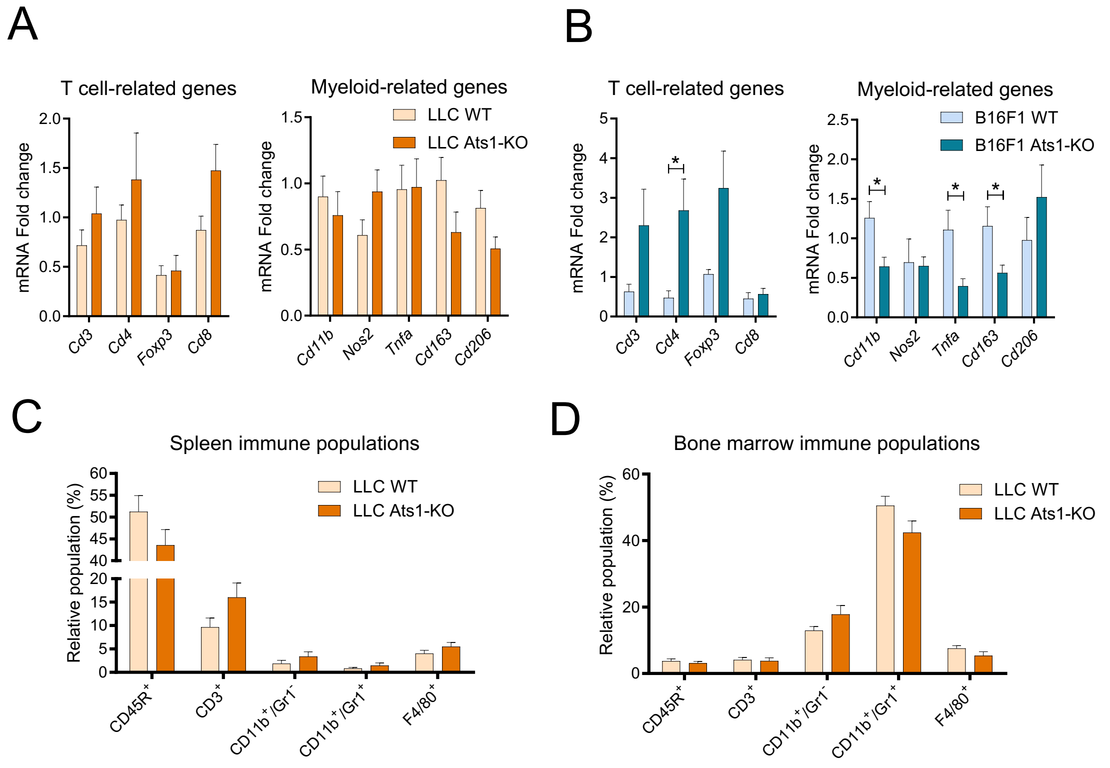

**Supplementary Figure 4. Comparison of myeloid cell populations in B16F1 WT and LLC WT tumours.** FC data showing the percentage of myeloid populations (CD11b$^+$/Gr1$^-$, CD11b$^+$/Gr1$^+$ and F4/80$^+$) represented as mean ± s.e.m of live cells in B16F1 WT and LLC WT tumours. (*, *p<0.05*; ***, *p<0.001*; two-tailed t Student, n=5 and n=9 samples in B16F1 WT and LLC WT models, respectively).

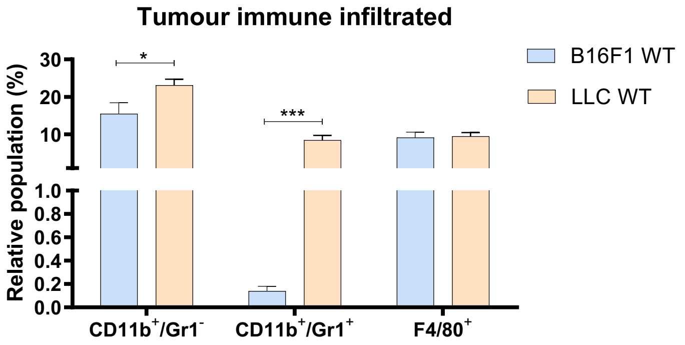

**Supplementary Figure 5. Functional characterisation of WT and Ats1-KO BMDMs.** (A) Representative gating strategy showing $CD11b^+$ population selection (mean percentage included) from bone marrow culture. (B) Representative bright field pictures of WT and Ats1-KO macrophages after selection. (C) Cell index representing the cell adhesion along the time performed in xCELLigence platform. (D) Representative immunofluorescence pictures of WT and Ats1-KO macrophages after 24 hours of polarisation to M1, M2 and non-polarised M0 BMDM. Cells were seeded and polarised on fibronectin-coated coverslips and stained with phalloidin-TRITC (red) and α-tubulin-FITC (green). White scale = 40μm, yellow scale = 20μm. (E) FC quantification of (up) NOS2 and (down) CD206 (represented as mean percentage ± s.e.m) in M1 and M2 24h-polarised $CD11b^+$ populations respectively (n=9 and n=4 samples in WT and Ats1-KO groups, respectively). (F) Representative pictures after 24-hour migration experiment in presence of FBS. Scale = 150μm. (G) Quantification of two independent experiments showing the relative migration of positive control (with FBS) compared to negative control (without FBS).

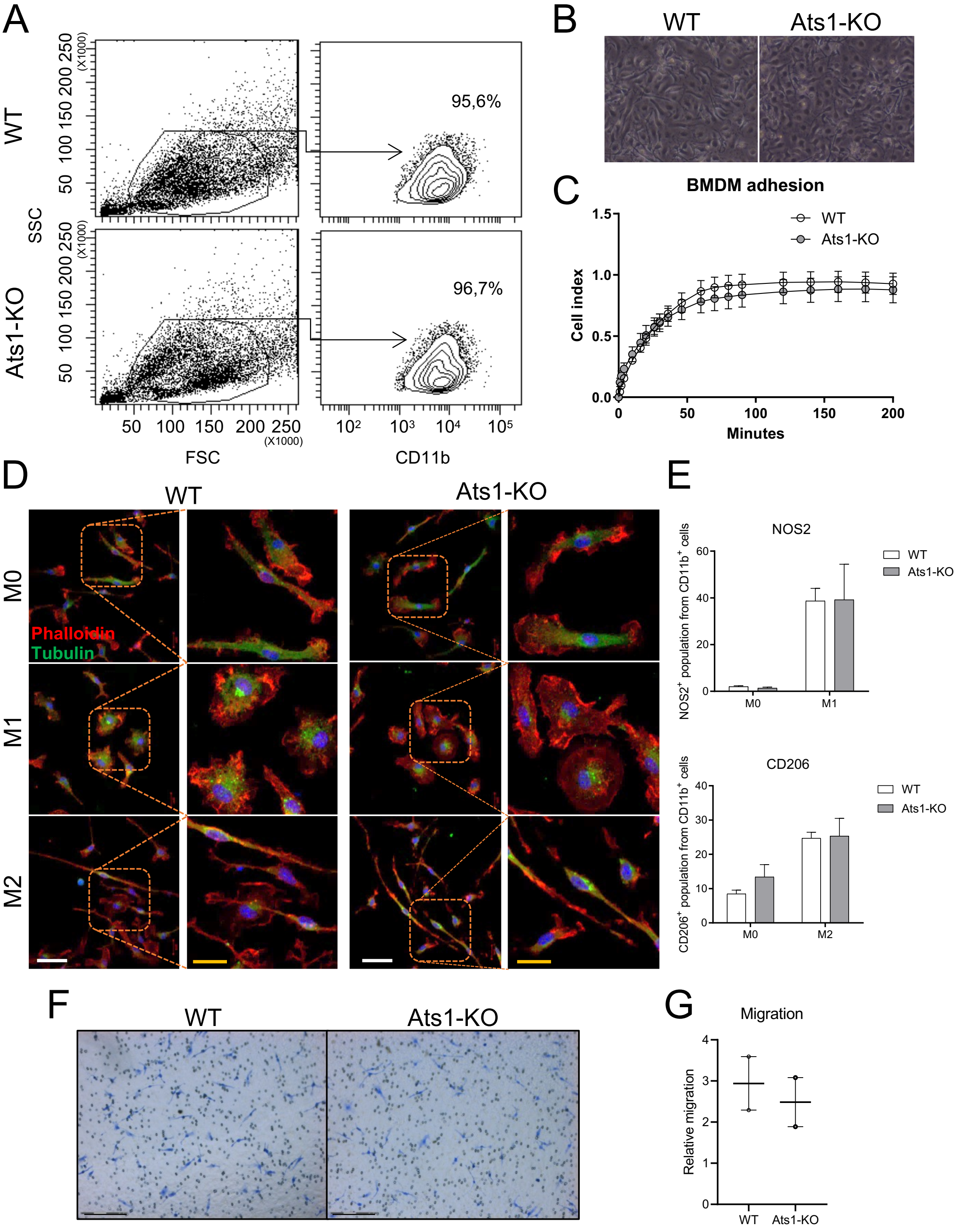
A
WT
Ats1-KO
SSC
FSC
CD11b
95,6%
96,7%
B
WT
Ats1-KO
C
BMDM adhesion
Cell index
Minutes
WT
Ats1-KO
D
WT
Ats1-KO
M0
M1
M2
Phalloidin
Tubulin
E
NOS2
NOS2+ population from CD11b+ cells
CD206
CD206+ population from CD11b+ cells
WT
Ats1-KO
F
WT
Ats1-KO
G
Migration
Relative migration
WT
Ats1-KO

**Supplementary Figure 6. Confirmation of Ats1-KO BMDM phagocytosis deficiency by clodronate liposomes *in vitro***

(A) Experimental schematic of the phagocytosis assay using clodronate liposomes *in vitro.* (B) Percentage of viable WT and Ats1-KO BMDM after clodronate treatment (represented as mean percentage ± s.e.m). (B) Percentage of apoptotic (Annexin V$^+$/7AAD$^-$) cells after clodronate treatment for both genotypes (represented as mean percentage ± s.e.m). Dot lines represent the treatment with PBS liposomes. N=3 samples in each group.

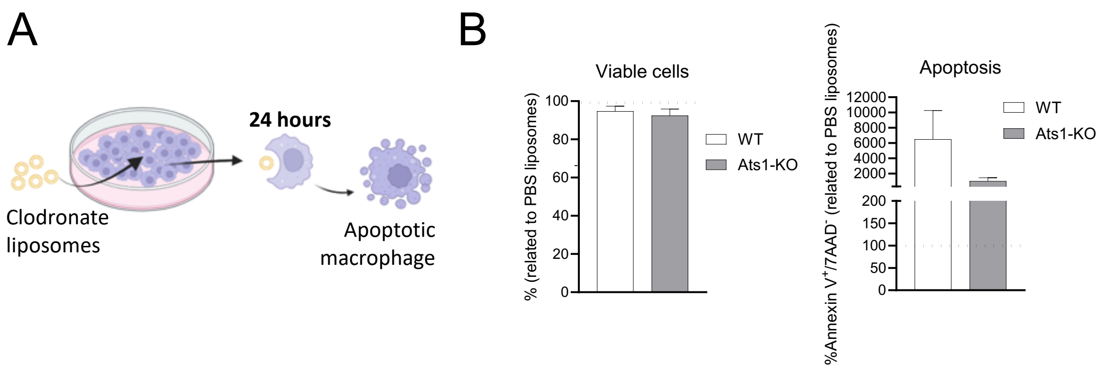
